# Co-targeting KRAS and Exportin1 as an effective therapeutic strategy for KRASG12D mutant pancreatic ductal adenocarcinoma

**DOI:** 10.1101/2025.11.21.689857

**Authors:** Husain Yar Khan, Mohammed Najeeb Al Hallak, Amro Aboukameel, Sahar F. Bannoura, Md. Hafiz Uddin, Adeeb A. Aboukameel, Fulya Koksalar Alkan, Ahmet B. Caglayan, Hilmi K. Alkan, Bin Bao, Hugo Jimenez, Allan M. Johansen, Callum McGrath, Grayson Barker, Khalil Choucair, Miguel Tubon, Eliza Beal, Steve Kim, Rafic Beydoun, Gregory Dyson, Yang Shi, Misako Nagasaka, Azeddine Atfi, Hasan Korkaya, Muhammad Wasif Saif, Philip A. Philip, Bassel El-Rayes, Herbert Chen, Anthony F. Shields, Ramzi M. Mohammad, Boris C. Pasche, Asfar S. Azmi

## Abstract

**Background:** Several KRASG12D inhibitors (KRASG12Di) are under clinical evaluation for pancreatic ductal adenocarcinoma (PDAC). However, as seen with other first generation KRAS inhibitors, resistance may limit their long-term efficacy, necessitating combination strategies to enhance therapeutic outcomes. Exportin 1 (XPO1), a nuclear transport protein overexpressed in PDAC, represents a therapeutic vulnerability in KRAS-mutant cancers. Here, we demonstrate that the second-generation XPO1 inhibitor Eltanexor synergizes with MRTX1133 to enhance its efficacy in multiple PDAC models.

**Methods:** We generated KRASG12Di-resistant PDAC cells and assessed their response to Eltanexor. The antiproliferative effects of MRTX1133 and Eltanexor combinations were evaluated in 2D and 3D *in vitro* PDAC models. The *in vivo* efficacy of the combination was tested in KRASG12D-mutant human and murine PDAC xenograft and allograft models.

**Results:** Eltanexor sensitized MRTX1133-resistant PDAC cells to growth inhibition. In both 2D and 3D culture models, the combination of Eltanexor and MRTX1133 significantly reduced cell viability. Mechanistically, the combination treatment suppressed key KRAS downstream signaling molecules, including p-ERK, mTOR, p-4EBP1, DUSP6, and cyclin D1. Kinome analysis further revealed reduced MAPK-related kinase activity. Combining subtherapeutic doses of Eltanexor and MRTX1133 resulted in significant tumor regression and prolonged survival in PDAC xenograft and immunocompetent orthotopic allograft models. Moreover, maintenance therapy with Eltanexor prevented tumor relapse, yielding a durable antitumor response.

**Conclusion:** This study demonstrates that Eltanexor overcomes resistance to MRTX1133 and enhances its efficacy in PDAC. The combination regimen may provide a durable therapeutic response while reducing the required dose of KRASG12D inhibitors, potentially delaying resistance and improving patient outcomes.

**Statement of Translational Relevance:** PDAC remains one of the deadliest malignancies, with limited effective therapies and dismal survival rates. The emergence of KRASG12D-selective inhibitors, such as MRTX1133, marks a critical advance for nearly 40% of PDAC patients harboring this oncogenic driver. However, inevitable emergence of adaptive or acquired resistance to KRAS inhibitors remains a major barrier to achieving durable clinical benefit. This study uncovers XPO1 inhibition as a rational and synergistic strategy to augment the antitumor efficacy of MRTX1133. By enhancing KRASG12D inhibitor activity and potentially reducing the required therapeutic dose, this combination approach offers a novel means to delay or overcome resistance. These findings provide a strong preclinical rationale for clinical trials evaluating KRAS inhibitors in combination with XPO1 inhibitors and may significantly improve outcomes for a substantial subset of PDAC patients who currently lack effective targeted treatment options.

## INTRODUCTION

Pancreatic cancer, which includes PDAC, is an aggressive malignancy with an exceptionally poor prognosis. Despite 481 phase 1 and 85 phase 3 trials since the year 2000, and five drug approvals, median survival in metastatic pancreatic cancer remains less than one year [1]. There has been a rise in 5-year survival for pancreatic cancer - from 4% to 13%, but it is largely due to increased detection of indolent neuroendocrine tumors; for PDAC, 5-year survival remains only 8% [2]. In addition to dismal survival rate, the lack of effective therapies further exacerbates the clinical burden of this disease that is characterized by a grim prognosis. PDAC remains a formidable challenge due to its late-stage diagnosis, intrinsic resistance to conventional chemotherapy and radiotherapy, and the ineffectiveness of immunotherapy. Consequently, there exists a critical want for novel therapeutic approaches capable of substantially improving outcomes for PDAC patients [3].

Oncogenic KRAS mutations occur in more than 90% of patients with PDAC [4]. A recent retrospective study of 803 PDAC patients found that patients with KRAS mutations, particularly G12D, had significantly shorter overall survival compared to those with wild type KRAS (22 vs. 38 months, *P* < 0.001) [5]. Moreover, KRASG12D mutations were significantly enriched in metastatic tumors compared to primary tumors (34% vs. 24%, *P* = 0.001) [5]. These observations indicate that KRAS status and subtype are associated with worse prognosis in PDAC.

The clinical development of KRAS mutation specific inhibitors (KRASi) has been a groundbreaking advancement in oncology. Despite showing promise as potent targeted therapies, KRASi as monotherapies have limited efficacy. Patients treated with these agents have short duration of response and develop drug resistance over time [6, 7], highlighting the need for combination approaches that can potentially enhance the sensitivity of tumors to KRASi when co-targeted.

Novel KRAS inhibitors that target the G12D isoform, a major KRAS mutation in PDAC, have been developed and are expected to benefit a large PDAC patient population [8]. However, learning from clinical experience with sotorasib and adagrasib (KRASG12Ci), there is a high possibility that the KRASG12Di as monotherapy will also show only a modest increase in disease-free survival as both G12D and G12C inhibitors mediate their effects by targeting essentially the same downstream signaling. Therefore, there is a pressing need to develop novel combination therapies that can enhance the efficacy and maximize the therapeutic potential of KRASG12D targeted drugs in PDAC.

The nuclear export protein Exportin 1 (XPO1) plays a vital role in maintaining cellular homeostasis by mediating the transport of various protein cargoes [tumor suppressor proteins (TSPs), genome surveillance proteins, transcription factors] and RNA molecules out of the cell nucleus [9]. Increased expression of XPO1 has been observed in PDAC which reportedly correlated with poor prognosis [10]. XPO1 overexpression enhances the export of TSPs to the cytosol, thereby preventing them from carrying out their normal function of cell growth regulation in the nucleus [11]. Therefore, XPO1 inhibition has emerged as an appealing anticancer strategy [9], especially in view of multiple reports implicating XPO1 to be a general vulnerability across several types of cancer [12]. Interestingly, it has been reported that KRAS-mutant cancer cells are dependent on XPO1 mediated nuclear export, rendering XPO1 a druggable vulnerability in KRAS-mutant cancer [13].

Earlier, we developed KRASG12Ci-resistant PDAC and NSCLC models which were found to be acutely sensitive to XPO1i Selinexor [14]. The combination of sub-MTD doses of Sotorasib and Selinexor could lead to superior inhibition of tumor growth compared to single agent treatments [14]. These findings imply that the inhibition of XPO1 can be a plausible therapeutic strategy to amplify the efficacy of KRASi and for mitigating resistance to KRASi. In view of the synthetic lethal interaction between XPO1 and KRAS [14], here we have tested a novel combination of KRASG12Di MRTX1133 with XPO1i Eltanexor and demonstrated its potential as an effective and more durable PDAC therapy. Eltanexor, also known as KPT8602, is an investigational, second-generation selective inhibitor of nuclear export with better tolerability than Selinexor [15].

Here we tested XPO1 inhibitor in combination with KRASG12D inhibitor in multiple PDAC models. Using a range of KRASG12D-mutant *in vitro* and *in vivo* preclinical models of PDAC, we demonstrate enhanced anticancer activity of Eltanexor and KRASG12Di combinations. Our results indicate that this novel combination therapy can potentially improve treatment outcomes in KRASG12D-mutant PDAC.

## MATERIALS AND METHODS

### Cell lines, drugs, and reagents

AsPC-1, HPAC, HPAF-II and Panc-1 cell lines were purchased from American Type Culture Collection (ATCC, Manassas, VA, USA). 6694c2 cells were purchased from Kerafast, while KPC-313 was developed from KPC mouse tumor in our lab. AsPC-1 was maintained in RPMI-1640 (ATCC, Manassas, VA, USA) and all the other cell lines were maintained in DMEM (Thermo Fisher Scientific, Waltham, MA, USA), supplemented with 10% fetal bovine serum (FBS), 100 U/mL penicillin, and 100 μg/mL streptomycin in a 5% CO_2_ atmosphere at 37 °C. All cell lines were authenticated by short tandem repeat (STR) profiling using the PowerPlex® 16 System (Promega, Madison, WI) at the Applied Genomics Technology Center, Wayne State University. Routine mycoplasma testing was performed using a PCR-based assay. Experiments were conducted within 20 passages of each cell line. MRTX1133 and Eltanexor were obtained from Selleck Chemicals (Houston, TX) and prepared as 10 mM stock solutions in DMSO. For all in vitro assays, the vehicle control consisted of culture medium containing 0.1% DMSO.

### Transient knockdown of XPO1

HPAF-II cells were seeded in 6-well plate and incubated overnight to reach 90% confluence the next day. The cells were then transfected with siRNA for XPO1 knockdown using lipofectamine 3000 (Life Technologies, Carlsbad, CA, USA) according to manufacturer’s instructions, diluted in serum-free DMEM media. Lipofectamine-siRNA complexes were added to the wells, and fresh media containing the drugs was added 24 hours later.

### Developing KRASG12Di (MRTX1133) resistant cell line

Previously, we successfully developed multiple KRASG12Ci-resistant cell lines [14]. Using a similar approach, we also generated KRASG12Di- and pan-RASi-resistant cellular model for this study. Briefly, KRASG12D-mutant human PDAC cell line, AsPC-1, was maintained in long term cell culture exposed to incremental doses of MRTX1133 or RMC-6236 to develop drug resistance. After approximately four months of continuous exposure to MRTX1133 or RMC-6236, resistant cell populations were established and designated AsPC1-MRTX1133-R or AsPC1-RMC6236-R, respectively. These cells were subsequently treated with varying concentrations of MRTX1133 or RMC-6236, and cell viability was assessed using the MTT assay. Drug resistance was determined by calculating the fold change in IC50 values between the resistant and parental (drug-naive) AsPC1 cells.

### Cell viability assay and synergy analysis

Cells were seeded in 96-well plates at a density of 3 × 10³ cells per well and incubated overnight. The following day, the culture medium was replaced with 100 µL of fresh medium containing serial dilutions of the respective drugs, prepared from stock solutions using the OT-2 liquid handling robot (Opentrons, Queens, NY, USA). After 72 hours of drug exposure, cell viability was assessed using the MTT (3-(4,5-dimethylthiazol-2-yl)-2,5-diphenyltetrazolium bromide) assay as previously described [16]. IC50 values were determined from six replicates per dose using GraphPad Prism 4 software.

For synergy analysis, cells were treated for 72 hours with varying concentrations of MRTX1133, Eltanexor, or their combination at a fixed drug ratio (six replicates per treatment). Cell viability was measured by MTT assay, and the resulting data were used to generate isobolograms and calculate combination index (CI) values using CalcuSyn software (Biosoft, Cambridge, UK).

### Colony formation assay

Cells were seeded at a density of 100 cells per well in 24-well plates or 500 cells per well in 6-well plates and treated with either single agents or drug combinations for 72 hours. Following treatment, the drug-containing medium was replaced with fresh growth medium, and the cells were incubated for an additional 7 days to allow colony formation. At the end of the incubation period, the medium was removed, and colonies were fixed with methanol and stained with crystal violet for 15 minutes. Plates were then washed, air-dried, and imaged to document colony formation.

### Spheroid formation and 3D viability assay

HPAF-II, HPAC, KPC-313 and 6694c2 cells were collected as single cell suspensions using cell strainer and resuspended in 3D Tumorsphere Medium XF (PromoCell, Heidelberg, Germany). 200 cells were plated in each well of ultra-low attachment 96-well plates (Corning, Durham, NC, USA). Spheroids growing in spheroid formation medium were exposed to either Eltanexor, or MRTX1133, or a combination of Eltanexor with MRTX1133 twice a week for one week (four replicates for each treatment). At the end of the treatment, spheroid images were captured under an inverted microscope and 3D CellTiter-Glo assay (Promega, WI) was performed according to the manufacturer’s protocol.

### Immunoblotting

For total protein extraction, cancer cells were lysed in RIPA buffer and protein concentrations were measured using the bicinchoninic acid (BCA) protein assay (PIERCE, Rockford, IL, USA). A total of 40 μg of protein lysate from treated or untreated cells was resolved using 4-20% mini-Protean TGX (#4561093; BioRad, Hercules, CA, USA) gradient gels and transferred onto nitrocellulose membranes according to standard procedure. The membranes were incubated with the following primary antibodies (Cell Signaling Technology, Danvers, MA, USA) at 1:1000 dilution in 3% BSA: anti-XPO1 (#46249), anti-phospho-ERK ½ (#4370), anti-ERK ½ (#9102), anti-DUSP6 (#50945), anti-phopho-4EBP1 (#2855), anti-mTOR (#2972), anti-PARP (#9532), anti-RB1 (#9309), anti-CDK4 (#12790), and anti-Cyclin D1 (#55506). While anti-GAPDH (#sc-47724; Santa Cruz Biotechnology, Santa Cruz, CA, USA) was used at a dilution of 1:3000. Incubation with 1:10000 diluted IRDye 800CW goat anti-mouse/IRDye 680RD goat anti-rabbit secondary antibodies (#827-08364/926-68171; LI-COR Biosciences, Lincoln, NE, USA) in 3% BSA solution was subsequently performed at room temperature for 1 hour. The signal was detected using the LI-COR Odyssey DLx Imager (Lincoln, NE, USA).

### Phosphokinome Profiling

Kinome profiling was performed using PamGene’s PamChip® technology (PamGene International BV, ’s-Hertogenbosch, Netherlands), which enables real-time measurement of global kinase activity to assess signaling alterations in cancer cells following drug treatment. HPAC cells seeded in 6-well plates (1 × 10^6^ cells per well) were treated with MRTX1133 (10 nM), Eltanexor (250 nM), or a combination of both agents for 6 hours. At the end of treatment, cells were washed with cold PBS and scraped on ice to prepare cell lysates. Cell lysates (three biological replicates per treatment) were applied to PamChip® 4 microarrays containing either 196 protein tyrosine kinase (PTK) or 144 serine/threonine kinase (STK) phosphosites, composed of 13-amino acid peptides immobilized on a 3D porous ceramic membrane. During the assay, lysates were actively circulated through the porous arrays to ensure efficient contact between kinases and their respective peptide substrates. Active kinases phosphorylated the immobilized phosphosites, which were detected using fluorescently labeled antibodies. Fluorescent signals were captured in real-time at multiple exposure times using a CCD camera integrated into the PamStation®. Image quantification, signal normalization, and statistical analysis were performed using PamGene’s BioNavigator® software. Differential phosphorylation patterns were used to infer upstream kinase activity and pathway modulation resulting from individual or combined treatment with MRTX1133 and Eltanexor.

### RNA isolation and RT-qPCR

Total RNA from HPAC cells were extracted and purified using the RNeasy Mini Kit and RNase-free DNase Set (QIAGEN, Valencia, CA) following the protocol provided by the manufacturer. The expression levels of MAP2K1, MAP2K2, MAP2K3, MAP3K5, MAP3K6, MST1R, ABL1, ABL2, ROS1 and SYK in single agent or combination treated HPAC cells were analyzed by real-time RT-qPCR using High-Capacity cDNA Reverse Transcription Kit and SYBR Green Master Mixture from Applied Bio-systems (Waltham, MA, USA). The conditions and procedure for RT-qPCR have been described previously [16]. Sequences of primers used are listed in **Table S1**.

### KRASG12D-mutant PDAC *in vivo* models

*In vivo* studies were conducted under Wayne State University’s Institutional Animal Care and Use Committee (IACUC) approved protocol in accordance with the approved guidelines. Experiments were approved by the institute’s IACUC (Protocol # 22-01-4355).

### Human PDAC cell-derived (AsPC-1 and HPAC) xenograft models

Post adaptation in our animal housing facility, 4-5 weeks old female ICR-SCID mice (Taconic Biosciences, Rensselaer, NY) were subcutaneously implanted with AsPC-1 or HPAC cells. 1×10^6^ cells suspended in 200 μL PBS were injected unilaterally into the left flank of donor mice using a BD 26Gx 5/8 1ml Sub-Q syringe. Once the tumors reached about 5-10% of the donor mice body weight, the donor mice were euthanized, tumors were harvested, and fragments were subsequently implanted into recipient mice. Seven days post-transplantation, the recipient mice were randomly divided into four groups of 6 mice each and received either vehicle, or Eltanexor (15 mg/Kg BIW, PO), or MRTX1133 (30 mg/Kg QD, IP), or their combination for four weeks. In addition to double randomization, to further reduce bias, blinding was observed during tumor measurement and data analysis. On completion of drug dosing, tumor tissue from control or treatment groups were harvested for immunohistochemical (IHC) analysis.

### Immunocompetent murine PDAC (6694c2) syngeneic allograft models

PDAC KPC 6694c2 cells were washed with PBS and then suspended in cold PBS at a concentration of 200,000 cells per 100 μL. The cell suspension was kept on ice and mixed with an equal volume of Matrigel matrix (# 356237; Corning, Durham, NC, USA). Using female C57BL/6 (Envigo, Indianapolis, IN, USA) mice, we performed subcutaneous double flank inoculation of 6694c2 cells mixed with ECM. Once tumor sizes reached ∼ 200 mm^3^, the mice were blindly randomized to receive vehicle, Eltanexor (15 mg/Kg BIW, PO), MRTX1133 (30 mg/Kg QD, IP) or a combination of both. For the survival study, surgical orthotopic implantation of murine PDAC 6694c2 cells (50,000 cells mixed with Matrigel in 1:1 ratio) in the head of the pancreas of female C57BL/6 mice was performed. Approximately two weeks post implantation when the tumors were palpable, mice were randomly divided into four treatment groups of seven mice each and treated blindly with either MRTX1133, or Eltanexor, or a combination of both at the aforementioned doses for four weeks.

### Immunostaining

Residual tumors were harvested immediately after euthanizing animals and were formalin-fixed for histopathology. Formalin-fixed tumor tissues were submitted to Karmanos Cancer Institute’s Histopathology Core for paraffin embedding. 5 μm thick serial sections were cut from formalin-fixed paraffin-embedded (FFPE) HPAC or 6694c2 tumor tissues and stained with hematoxylin-eosin (H&E) or IHC. The following antibodies (Cell Signaling Technology, Danvers, MA, USA) were used for IHC staining - anti-phosho-ERK1/2 (#4370) and anti-phosho-S6 (#2211) at 1:400 and 1:100 dilution, respectively.

### Statistical analysis

Wherever suitable, the experiments were performed at least three times. The data were also subjected to unpaired two-tailed Student *t* test wherever appropriate, and *P < 0.05* was considered statistically significant. For the kinome profiling, significance was obtained using one-way ANOVA followed by post-hoc Dunnett’s test (*P < 0.05*).

## RESULTS

### XPO1 inhibition enhances sensitivity to KRASG12Di and sensitizes KRASG12Di-resistant PDAC cells to growth inhibition

The co-expression analysis of 179 PDAC patient samples in TCGA show statistically robust correlation (Spearman’s correlation coefficient, ρ = 0.82; *P* = 2.08e-45) between KRAS and XPO1 (**Figure 1A**). A Spearman’s correlation coefficient value of 0.82 is indicative of a strong positive correlation (ρ > 0.5) between KRAS and XPO1, which means that as KRAS expression increases, XPO1 expression also tends to increase in these PDAC patients. To determine the functional role of XPO1 in KRASG12D-mutant PDAC, we knocked down the expression of XPO1 using RNA interference (siRNA) in HPAF-II human PDAC cell line (**Figure 1B**). The treatment of XPO1 knocked down (siXPO1) PDAC cells with KRASG12Di MRTX1133 at both low and high doses resulted in significantly more pronounced inhibition in cell viability compared to HPAF-II cells transfected with control siRNA, indicating that XPO1 inhibition could enhance the growth inhibitory activity of MRTX1133 in KRASG12D-mutant PDAC cells (**Figure 1C**).

**Figure 1:**
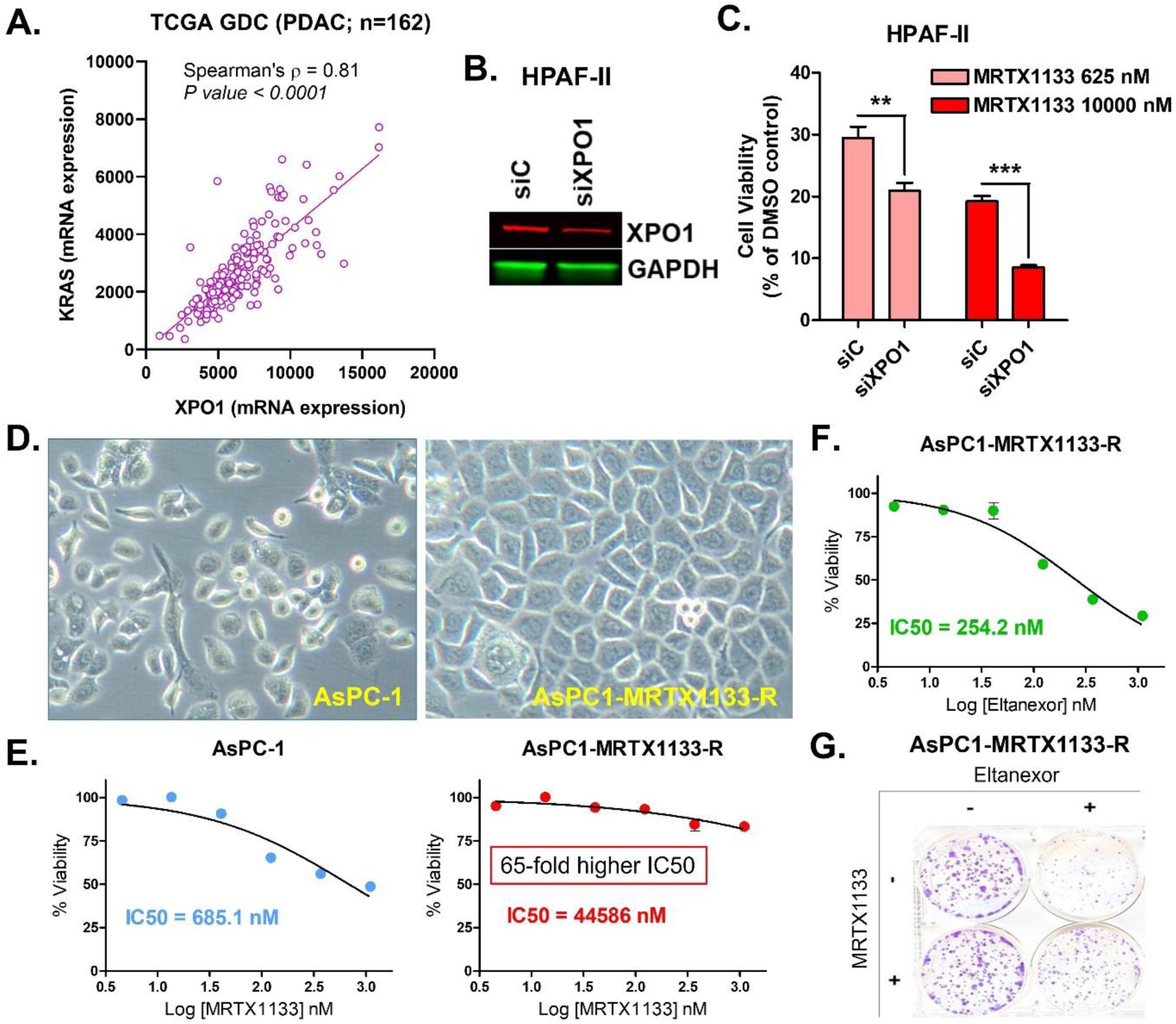
XPO1 inhibition enhances sensitivity to KRASG12Di and induces growth inhibition in PDAC cells resistant to KRASG12Di. **[A]** Spearmen correlation between KRAS and XPO1 in 179 patients with PDAC. **[B]** Western blot showing efficiency of siRNA mediated XPO1 silencing at 72 hrs. **[C]** Bar graph showing reduced cell viability in XPO1 knockdown HPAF-II cells treated with MRTX1133 compared to DMSO treated controls. Results are expressed as percentage of control ± S.E.M of six replicates. **[D]** MRTX1133 sensitive and resistant PDAC cells. AsPC1-MRTX1133-R cells show unresponsiveness to MRTX1133 **[E]** but exhibit sensitivity toward Eltanexor-induced growth inhibition **[F]** and inhibition of colony formation **[G]**. ** *P < 0.01*, *** *P < 0.001*

Furthermore, we generated KRASG12Di MRTX1133-resistant human PDAC cells (AsPC1-MRTX1133-R) *in vitro,* which appear morphologically distinct from the parental AsPC-1 cells indicting drug induced lineage plasticity (**Figure 1D**). AsPC1-MRTX1133-R cells exhibit more than 65-fold increase in IC50 value for MRTX1133 compared to the drug sensitive AsPC-1, confirming that they have developed resistance to KRASG12Di (**Figure 1E**). Subsequently, this MRTX1133-resistant cell line was treated with Eltanexor and was found to be sensitive to Eltanexor-induced cell growth inhibition (**Figure 1F**) and suppression of colony formation (**Figure 1G)**. This establishes that PDAC cells resistant to KRASG12Di respond to treatment with Eltanexor.

### Synergistic effects of KRASG12Di and XPO1i combination *in vitro*

We tested various dose combinations of KRASG12Di (MRTX1133) and XPO1i (Eltanexor) for their antiproliferative effects on PDAC cellular models in 2D culture. Both MRTX1133 and Eltanexor reduced the ability of the murine PDAC KPC-313 cells to form colonies in a dose-dependent manner (**Figure 2A**) and a combination of Eltanexor with MRTX1133 was strikingly more effective at suppressing the clonogenic potential of these cells (**Figure 2B**). The combination of MRTX1133 and Eltanexor markedly suppressed the long-term survival and proliferative capacity of multiple KRASG12D-mutant PDAC cell lines, as evidenced by a remarkable reduction in colony formation (**Figure 2C-E**). This result indicates a robust and durable antiproliferative effect of the combination. Further, treatment of a panel of KRASG12D-mutant human PDAC cell lines, namely HPAF-II, HPAC, AsPC-1, Panc-1 and murine KPC 6694c2, with Eltanexor and MRTX1133 at different dose combinations synergistically inhibited cell proliferation as indicated by the combination index (CI) values less than 1 (**Figure 2F-J, Figure S1**). CI values were generated by the CalcuSyn synergy analysis performed using growth inhibition data. CI < 1 is synergistic, while CI > 1 signifies antagonistic effect.

**Figure 2:**
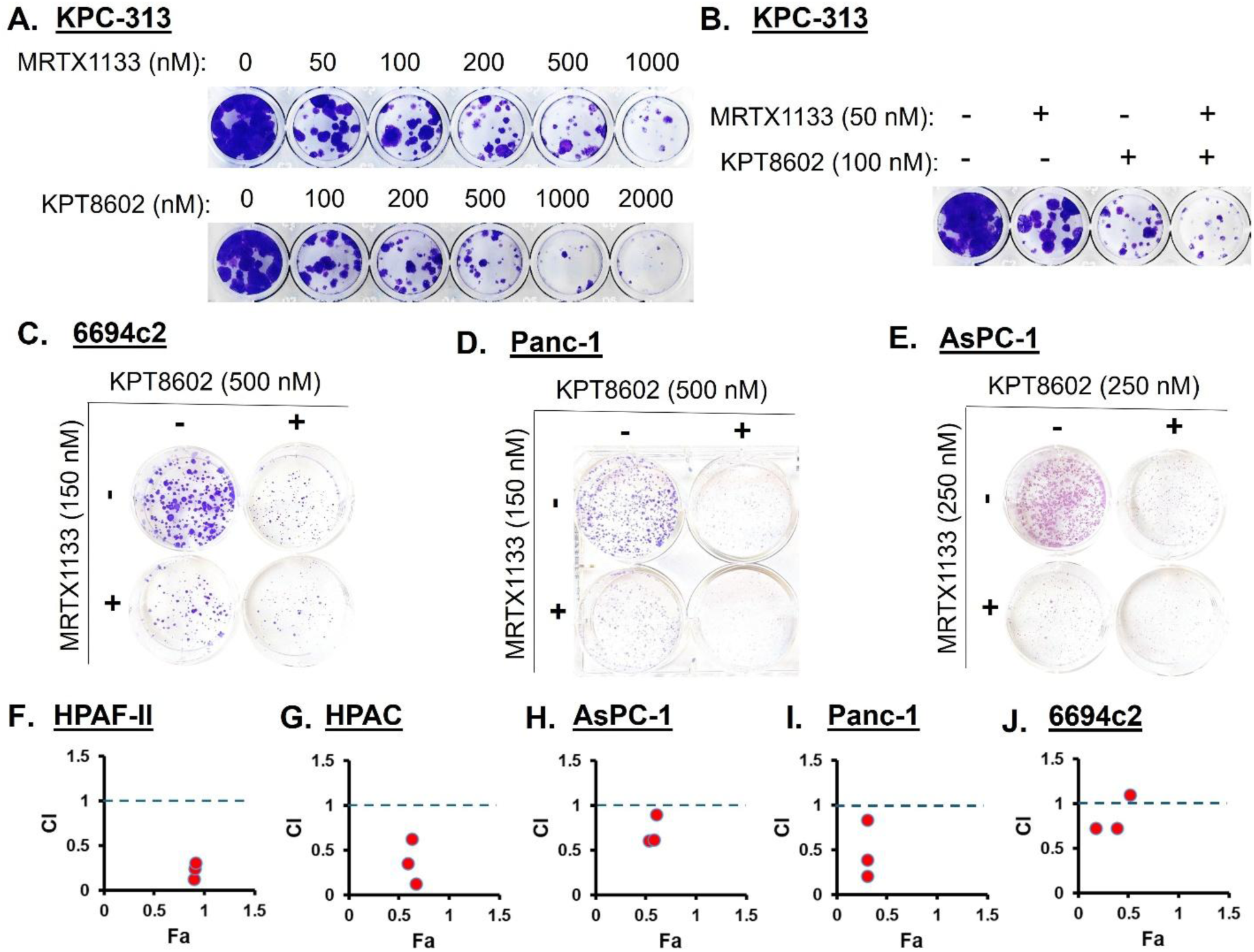
Synergistic effects of KRASG12Di and XPO1i combination *in vitro.* **[A]** Dose-dependent effects of MRTX1133 and Eltanexor, **[B]** as well as their combination on clonogenic potential of KPC-313 cells. **[C-E]** The combination suppressed the survival of multiple KRASG12D-mutant PDAC cell lines in colony formation assays. **[F-J]** Synergistic inhibitory effects of combination treatments on the growth of several PDAC cell lines in 2D culture in 72 hrs MTT assays. CalcuSyn software was employed to determine CI values from the resulting data. CI < 1 indicates synergistic effect of the drug combination at the corresponding doses. All results are expressed as percentage of control ± S.E.M of six replicates. Data is representative of three independent experiments.

### XPO1i and KRASG12Di combinations effectively disrupt the formation of KRASG12D-mutant PDAC cell-derived spheroids

Cell sensitivity in 3D culture is regarded as a more accurate predictor of *in vivo* efficacy, showing strong correlation with drug response in xenograft models [17]. Therefore, we performed a spheroid formation assay, where Eltanexor and MRTX1133 combination treatment resulted in reduced size and increased disintegration of spheroids derived from HPAF-II, HPAC and 6694c2 cell lines (**Figure 3A-C**). Further, exposing 3D cultures of multiple KRASG12D-mutant PDAC cells with combination of MRTX1133 and Eltanexor caused enhanced suppression of 3D cell viability (**Figure 3D-G**). Moreover, this effect of the combination treatment on 3D cell viability was found to be synergistic at multiple dose combinations tested as demonstrated by CI value < 1 (Figure 3H-J, Figure S2).

**Figure 3:**
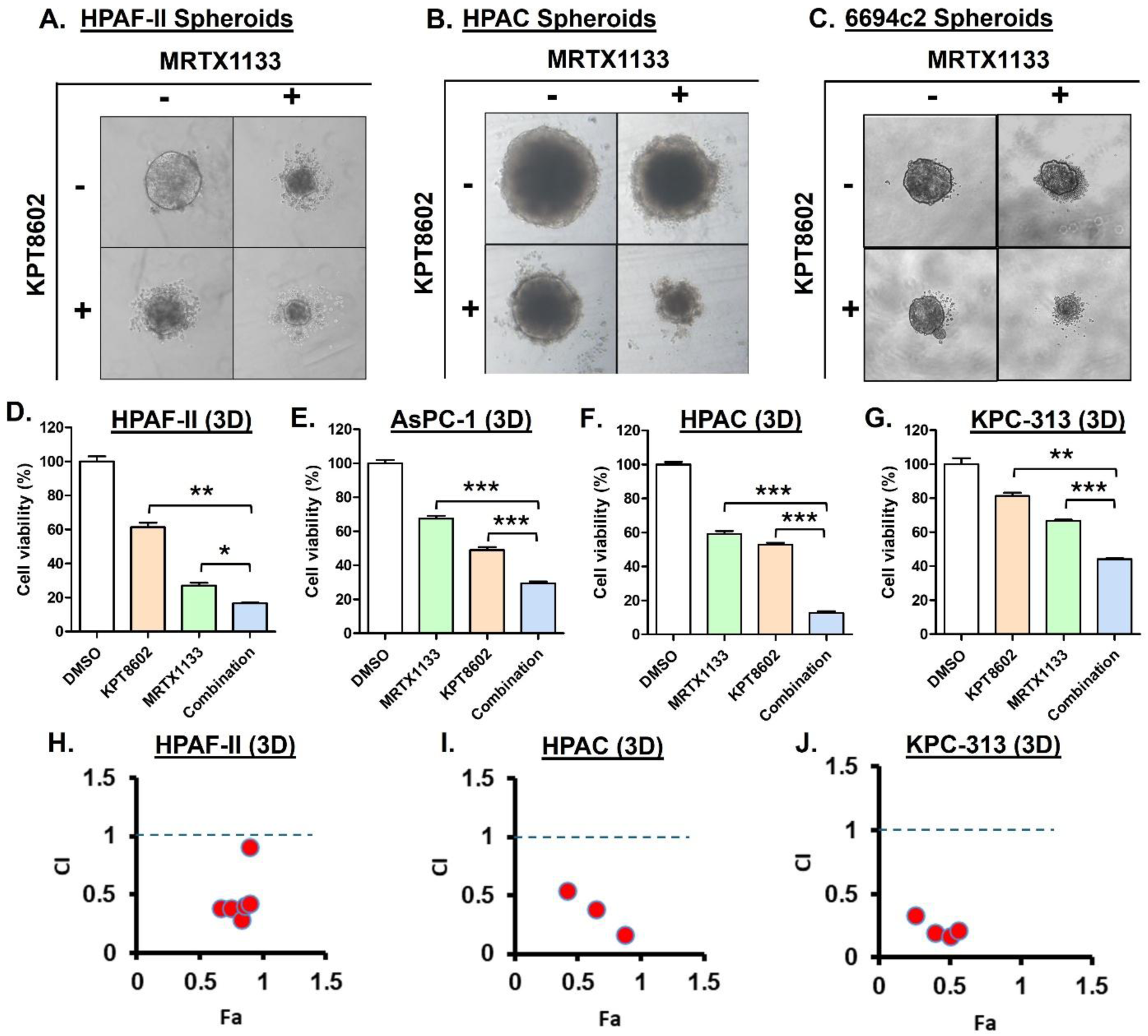
KRASG12Di synergizes with XPO1i to effectively disrupt the formation of KRASG12D mutant PDAC cell derived spheroids. **[A-C]** Combination of MRTX1133 (125 nM) and Eltanexor (125 nM) triggered enhanced suppression of spheroid formation. **[D-G]** The combination also induced enhanced cell viability inhibition of PDAC cells in 3D cultures as determined by 3D CellTiter-Glo assays performed one week post drug treatment. HPAF-II and AsPC-1 3D cultures were exposed to either Eltanexor or MRTX1133 or their combination at 125 nM. The concentration of MRTX1133 and Eltanexor used to treat HPAC and KPC-313 3D cultures was 250 nM each. **[H-J]** Synergistic inhibitory effects of the combination treatment at multiple doses of MRTX1133 and Eltanexor in constant ratio on PDAC cells in 3D cultures, as demonstrated by CI < 1 determined by CalcuSyn. All results are expressed as percentage of control ± S.E.M of four replicates. Data is representative of three independent experiments. * *P < 0.05*, ** *P < 0.01, *** P < 0.001*

### Combination of KRASG12Di and XPO1i modulate KRAS signaling and kinase activity

Next, we performed expression analysis to capture the impact of combination treatment on KRAS pathway molecules. Western blotting results show that the MRTX1133-Eltanexor combination treatment resulted in a reduction of the expression levels of p-ERK and its downstream DUSP6 as well as mTOR and its downstream p-4EBP1 in KRASG12D-mutant PDAC cells (**Figure 4A, Figure S3**). The combination also increased the expression of cleaved PARP, indicating enhanced apoptosis, as well as suppression of the tumor suppressor protein RB1, thereby preventing cell cycle progression through downregulation of cyclin D1 and CDK4 expression (**Figure 4B**).

**Figure 4:**
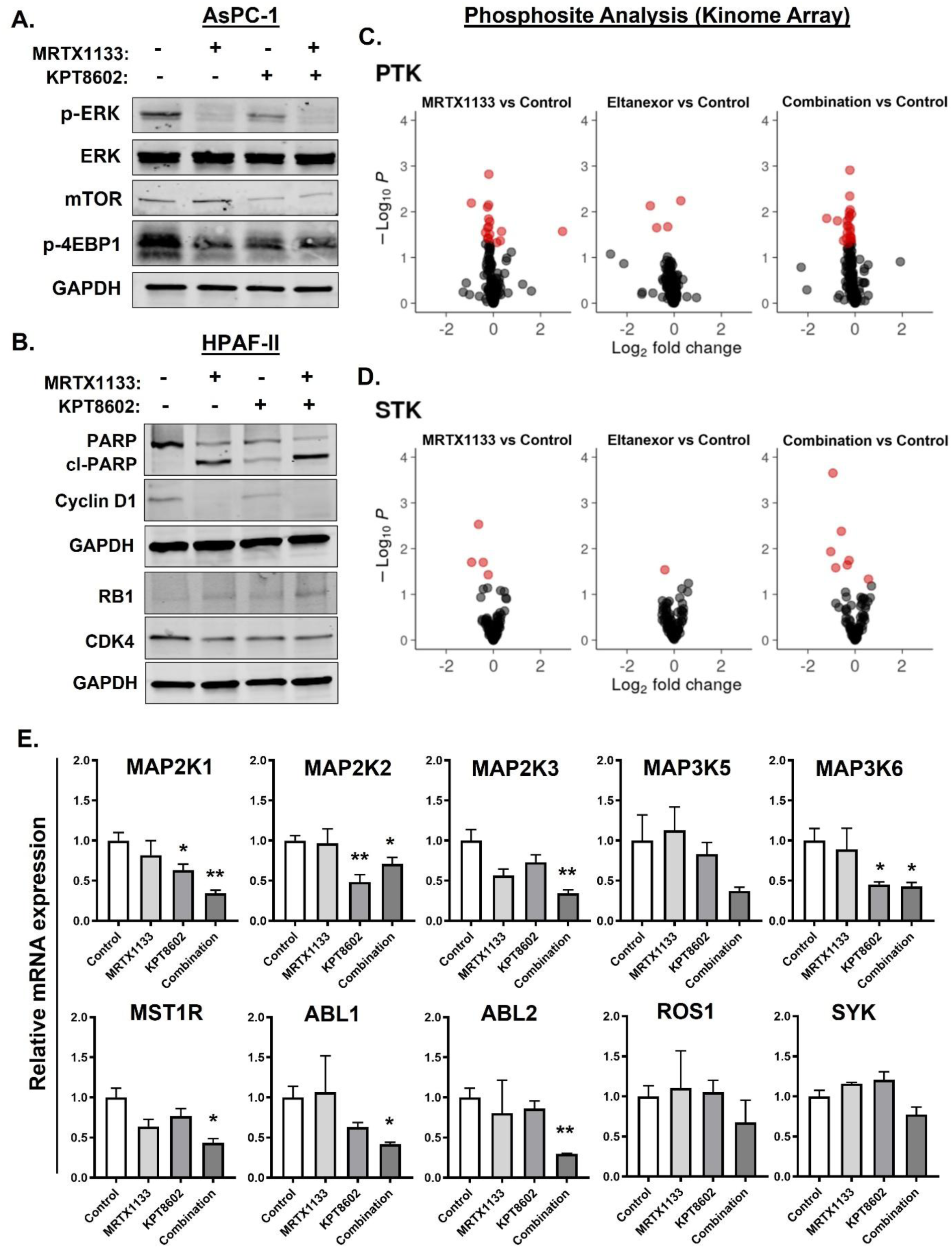
Combination of KRASG12Di and XPO1i modulates KRAS signaling and kinase activity. **[A-B]** Immunoblots showing reduced expression of key KRAS downstream signaling molecules and cell cycle markers as well as increased apoptosis marker cl-PARP in combination treated PDAC cells. AsPC-1 cells were treated with 250 nM and 500 nM of MRTX1133 and Eltanexor, respectively for 6 hrs. HPAF-II was exposed to 500 nM dose of either of the two drugs as single agents or in combination for 24 hrs. **[C-D]** Kinome profiling performed on MRTX1133 (10 nM) and Eltanexor (250 nM) treated HPAC cell lysates showing differentially phosphorylated PTK and STK phosphosites between conditions representing the overall trend of the treatment effect identified using ANOVA-Dunnett test. Red spots are phosphosites that show significant difference compared to untreated control (*P < 0.05*, i.e. -log10(*P*)>1.3). For each condition 3 biological replicates were analyzed. **[E]** Combination treatment induced alterations in mRNA expression were used to validate several of the identified kinases by RT-PCR. HPAC cells were treated with 10 nM MRTX1133 and 250 nM KPT8602 (Eltanexor) for 6 hrs before total RNA were isolated. Expression levels were normalized to β-actin and are presented as relative expressions compared to control. Data represents mean ± SEM of three replicates. * *P < 0.05*, ** *P < 0.01*

In addition, to better understand the underlying mechanisms of synergy and get mechanistic insights into signaling adaptations and potential vulnerabilities induced by the combination, we performed high throughput phospho-kinome profiling on the MRTX1133-Eltanexor treated PDAC cells. The kinome analysis can detect 196 PTKs and 144 STKs by measuring the phosphorylation of kinase substrates (phosphosites) immobilized on microarrays. Several of these kinases are relevant to tumor growth and are part of cell survival signaling. Alterations in the activity of PTKs and STKs in cell lysates of drug treated PDAC cells were evaluated. Our results indicate a higher number of significantly altered phosphosites (PTK: 24 vs 13; STK: 6 vs 4, P < 0.05) in the MRTX1133-Eltanexor combination compared to MRTX1133 alone treated cell lysates (**Figure 4C-D, Table S2**). Further, Upstream Kinase Analysis (UKA) showed clear reduction in kinase activity of several kinases related to MAPK signaling and multiple other kinases like MSTR1, ABL, ROS and SYK in the combination treated HPAC cell lysates (**Figure S4**).

Several of the identified kinases were validated by measuring changes in their mRNA expression levels upon drug treatment (**Figure 4E**). Notably, the combination treatment reduced the activity and expression of the cytosolic kinases MAP2K1 (MEK1) and MAP2K2 (MEK2) of the RAS-RAF-MEK-ERK pathway, which phosphorylate and activate ERK1/2, leading to downstream transcriptional changes that drive oncogenesis in KRAS-mutant PDAC. Combination treatment also downregulated the activity of the non-receptor tyrosine kinases ABL1 and ABL2 which regulate cytoskeletal remodeling, DNA damage response, apoptosis, and invasion/metastasis. Interestingly, both these kinases shuttle between the nucleus and cytoplasm. Of note, the two kinases regulate cell adhesion and motility in the cytoplasm, while they modulate DNA damage response and apoptosis in the nucleus [18].

### Preclinical antitumor efficacy of KRASG12Di and XPO1i combination in KRASG12D-mutant PDAC *in vivo* models

We employed KRASG12D-mutant human PDAC AsPC-1 cell-derived xenograft model to demonstrate superior efficacy of Eltanexor and MRTX1133 combination in suppressing tumor growth. In this model, treatment with MRTX1133 at an MTD dose (30 mg/Kg) resulted in tumor regression. Interestingly, the combination treatment caused significantly greater tumor regression compared to single agent MRTX1133 (**Figure 5A**). Notably, all animals in the combination cohort had tumor regression in contrast to only 3 out of 6 in the MRTX1133 alone group (**Figure 5B**). It is also worth mentioning that 4 animals (67% of the cohort) receiving combination treatment had >30% shrinkage in tumor volumes which may be considered analogous to a partial response (PR) observed in a clinical setting typically based on RECIST (Response Evaluation Criteria in Solid Tumors) guidelines. Moreover, both the drugs and their combination were tolerable as all the groups had < 10% body weight loss over the duration of the study (**Figure S5A**).

**Figure 5:**
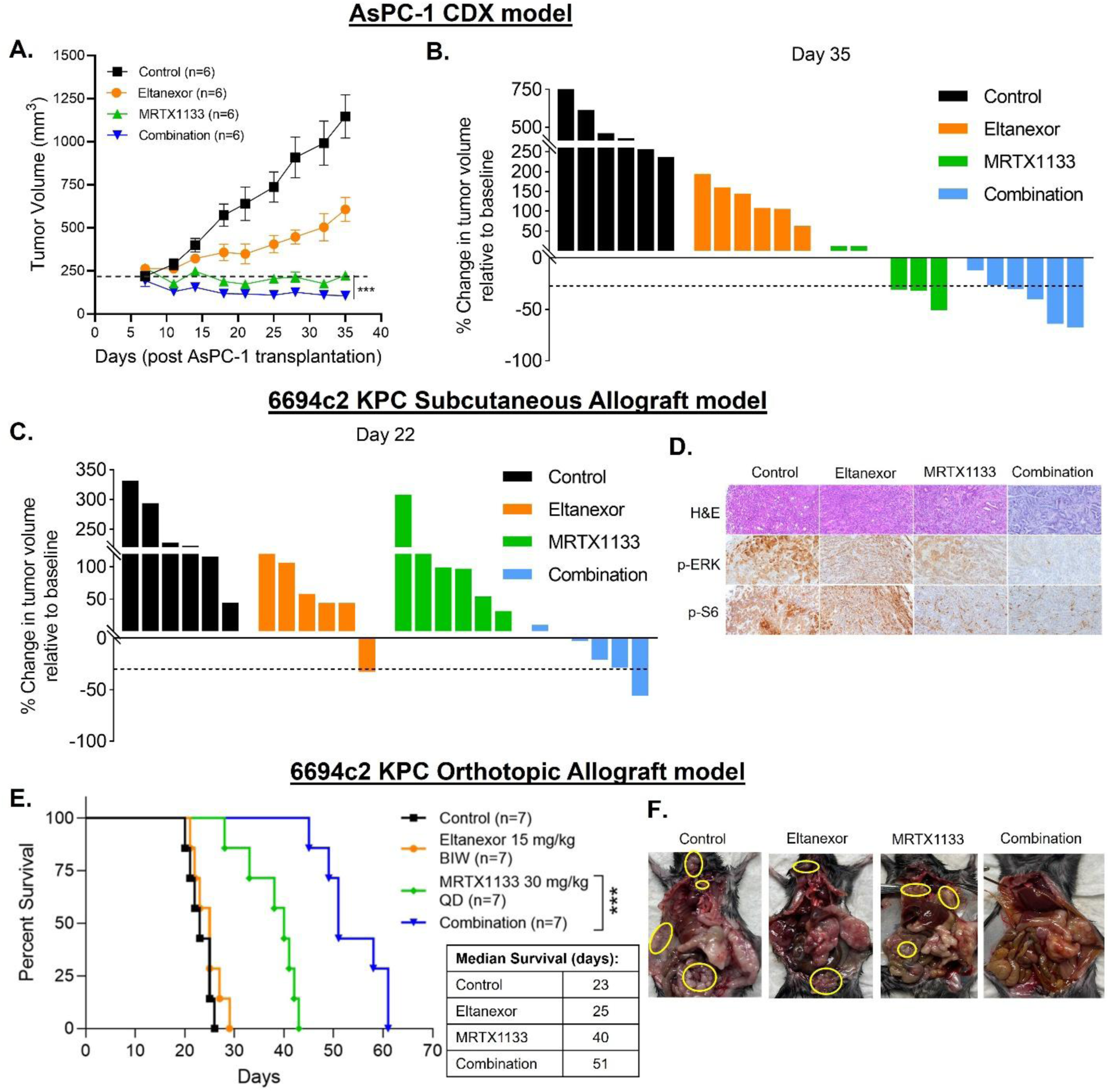
Preclinical antitumor efficacy of KRASG12Di and XPO1i combination. **[A-B]** The combination treatment results in superior tumor regression in ICR-SCID mice subcutaneously engrafted with AsPC-1 cells and administered MRTX1133 (30 mg/Kg QD, IP) and Eltanexor (15 mg/Kg BIW, PO) for 4 weeks. Tumor volumes in the combination group were determined to be statistically significant compared to those in the single agent MRTX1133 by a two-tailed unpaired Student’s *t*-test (*** *P < 0.001*). **[C]** Waterfall plot showing greater regression in 6694c2 KPC tumor volumes at the end of treatment (relative to baseline) in the combination treated C57BL/6 mice compared to either of the single agents alone. **[D]** IHC analysis showing (x100 magnification) reduction in p-ERK and p-S6 in tumor tissues harvested from mice treated with the combination of MRTX1133 and Eltanexor. **[E]** Kaplan-Meier plot showing significantly enhanced survival in combination treated cohort of an orthotopic PDAC allograft C57BL/6 mouse model. Statistical comparison of survival was performed by Log-rank (Mantel-Cox) test (*** *P < 0.001)*. **[F]** Distant metastasis can be seen in animals treated with single agent MRTX1133 or Eltanexor, while no metastasis is visible in mice treated with the combination.

In addition, we also tested the effect of MRTX1133-Eltanexor combination in an immunocompetent KRASG12D-mutant PDAC GEM (KPC) cell derived allograft (6694c2) model, where a remarkably superior antitumor efficacy of the combination was observed (**Figure 5C, Figure S5B**). Further, tumor tissue sections from the combination group showed reduced p-ERK and p-S6 expressions suggesting suppression of KRAS mediated cell growth signaling (**Figure 5D**). The combination treatment was also tested in an orthotopic murine PDAC syngeneic allograft (6694c2 KPC) model. As shown in **Figure 5E**, the median survival of animals in combination treated group was significantly better compared to those in the single agent cohorts. Furthermore, necropsy of mice treated with Eltanexor or MRTX1133 alone revealed distant metastatic nodules (**Figure 5F**). However, MRTX1133-Eltanexor combination treatment resulted in undetectable metastatic burden in this orthotopic immunocompetent PDAC mouse model demonstrating anti-metastatic effect of the combination **(Figure S5C)**. Therefore, these results establish that treatment with a combination of KRASG12Di and XPO1i leads to significantly improved control of primary tumor growth as well as metastatic spread compared to either treatment alone.

### Eltanexor treatment enhances the durability of response to KRASG12Di

To evaluate the therapeutic durability of Eltanexor in combination with KRASG12D inhibition, we utilized an HPAC-derived xenograft model. Eltanexor was administered orally at 15 mg/kg twice weekly throughout the study, while MRTX1133 was given at a sub-MTD dose of 10 mg/kg twice daily until day 35 post-engraftment (**Figure 6A**). During the initial treatment phase, both the combination and MRTX1133 monotherapy groups exhibited comparable tumor growth suppression. However, upon discontinuation of MRTX1133 at day 35, tumors in the MRTX1133 monotherapy group rapidly relapsed, indicating a lack of durable control. In contrast, mice in the combination group that continued receiving Eltanexor maintained tumor remission, suggesting that Eltanexor effectively suppressed tumor regrowth after KRASG12Di withdrawal. Strikingly, when MRTX1133 was reintroduced at day 50, it failed to re-establish tumor control in the monotherapy group, indicating the emergence of treatment resistance. Conversely, reinitiating MRTX1133 in mice that had remained on Eltanexor led to a sustained and durable response (**Figures 6B and 6C**). Importantly, all treatment arms were well tolerated, with <10% body weight loss over the study period (**Figure 6D**). Immunohistochemical analysis of tumor tissues from the combination-treated cohort revealed a marked reduction in p-ERK expression, confirming suppression of downstream KRAS signaling (**Figure 6E**). These findings demonstrate that Eltanexor can significantly extend the durability of response by preventing relapse and possibly overcoming acquired resistance.

**Figure 6:**
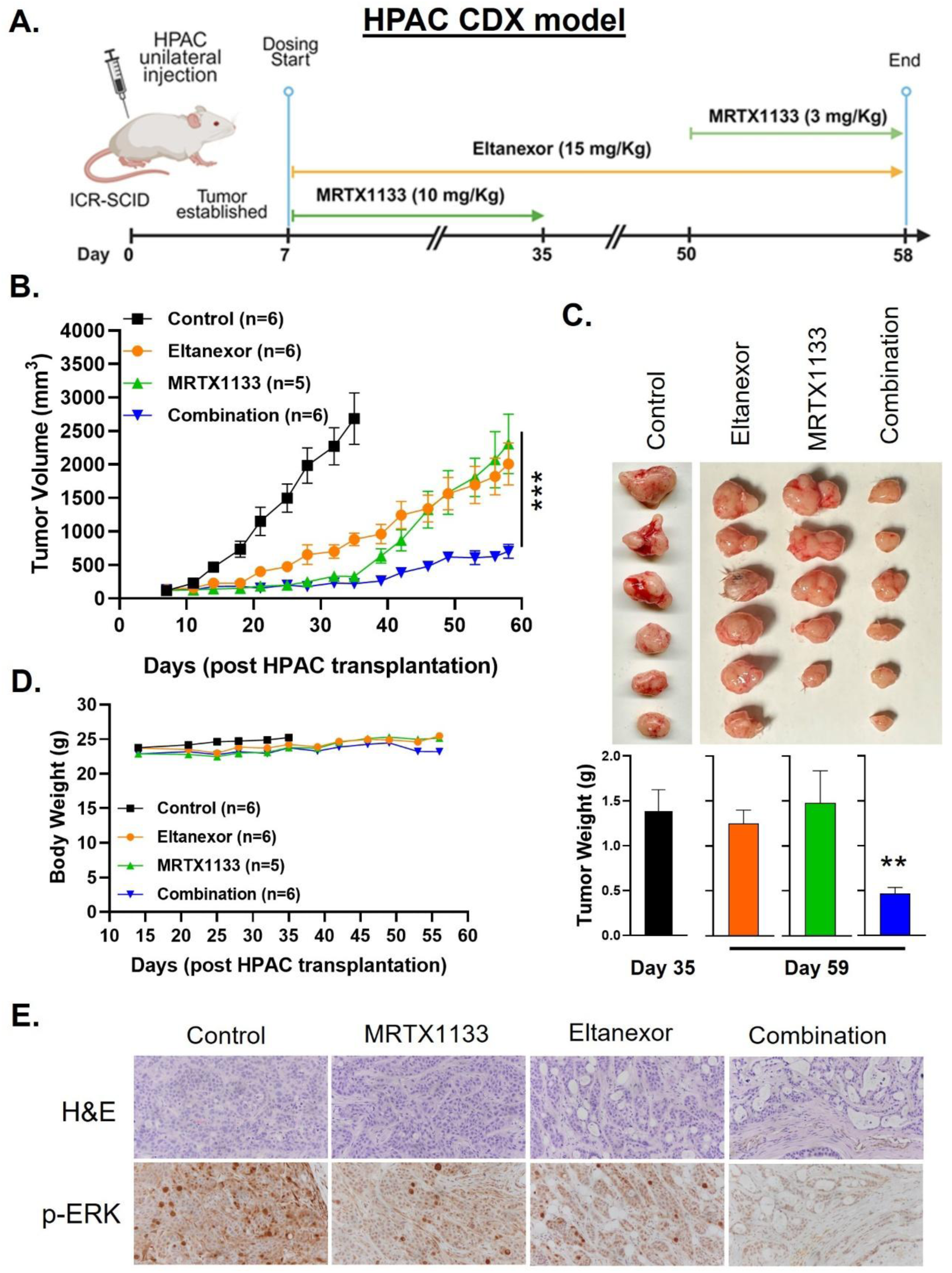
Maintenance with Eltanexor in combination treated mice prevents tumor relapse and results in a durable response. ICR-SCID mice subcutaneously engrafted with KRASG12D-mutant PDAC (HPAC) cells were intraperitoneally administered MRTX1133 at 10 mg/Kg BID until day 35. MRTX1133 treatment was stopped for two weeks while Eltanexor was maintained throughout the course of the study at an oral dose of 15 mg/Kg BIW. Mice were readministered MRTX1133 starting on day 50 albeit at a lower dose of 3 mg/Kg. **[A]** timeline for the subcutaneous HPAC CDX model. This study timeline has been created with BioRender.com (License # *QZ286IJ3IM*). Drug treatment induced changes in **[B]** tumor volumes, **[C]** tumor weights, **[D]** body weights, and **[E]** p-ERK expression in residual tumor tissues. ** *P < 0.001*, *** *P < 0.001*

## DISCUSSION

In this article, we for the first time show synergy between KRASG12D inhibitor MRTX1133 and nuclear export protein XPO1 inhibitor Eltanexor. Our combination approach of co-targeting KRASG12D and XPO1 resulted in enhanced growth suppression of KRASG12D-mutant PDAC cells and cell-derived tumor xenografts. This study brings forward a novel combination therapy for drug resistant KRASG12D-mutant tumors and provides preclinical rationale for the use of Eltanexor in a clinical setting to prevent or delay the development of resistance in patients receiving KRASG12D inhibitor monotherapy.

KRAS mutations are a key driver of oncogenic alterations in various cancers, particularly in PDAC. As one of the most frequently mutated oncogenes, KRAS was long deemed undruggable, posing a significant challenge for targeted cancer therapy [19]. However, a breakthrough came a decade ago when molecules that can bind to the G12C isoform of KRAS protein were developed [20]. This revived the quest for targeting KRAS which ultimately resulted in FDA approval in 2021 for the first KRASi sotorasib for the treatment of NSCLC patients carrying KRASG12C mutation [21], followed by approval for the second KRASi adagrasib the very next year [22]. Despite advances in targeting KRAS oncoprotein achieved by the clinical development of KRASG12Ci, PDAC patients are miles away from being benefitted by such promising therapies. Although KRAS mutations are present in 93% of PDAC patients, KRASG12C accounts for less than 2% of them. The most common KRAS mutation in PDAC is KRASG12D (42% of patients) [23]. Compared to other KRAS mutations, the G12D mutation has been reported to exhibit the highest oncogenic potential in preclinical models of PDAC [24]. Moreover, PDAC patients carrying the KRASG12D mutation show a shorter survival duration compared to those with wild-type KRAS or other major KRAS mutations [25].

Recently, several KRASG12D inhibitors have been developed and are in phase I clinical evaluation. MRTX1133 is a small molecule selective inhibitor of KRASG12D that has been developed through extensive structure-based drug design. MRTX1133 has shown potent *in vitro* and *in vivo* antitumor effects against KRASG12D-mutant PDAC models [26], leading to its clearance as an investigational new drug by the FDA for a phase I/II clinical trial (NCT05737706), treating patients with advanced solid tumors including PDAC. More recently, pan-RAS inhibitors (RMC-6236 and RMC-7977) have been developed and are currently under clinical evaluation as well [27]. Nevertheless, targeting KRAS in pancreatic cancer remains challenging. While MRTX1133 and other KRAS inhibitors show potential to become promising therapy for PDAC patients harboring KRASG12D mutations, their impact on clinical management of PDAC may be limited without complementary therapeutic strategies.

It is noteworthy that PDAC patients receiving monotherapy of KRASi sotorasib and adagrasib have a modest objective response rate (ORR) of 21% and 33.3%, respectively [6, 7]. Similar to other KRAS targeted drugs, KRASG12Di also have limited efficacy as monotherapy and resistance ultimately develops in most patients, which impedes their prolonged therapeutic use. This necessitates the identification of combination therapies that can bolster the efficacy of KRASG12Di and either prevent the emergence or at least delay the onset of drug resistance in PDAC patients receiving KRASG12Di. In this regard, we aim to develop a novel combination therapy that can enhance the efficacy of KRASG12Di, which in turn can reduce the effective dose of KRASi thereby circumventing or delaying therapeutic resistance and achieving a durable antitumor response.

There is growing interest in identifying KRAS-associated synthetic lethal interactions and developing small molecule inhibitors targeting these vulnerabilities. In a comprehensive multi-genomic analysis of 106 human NSCLC cell lines, Kim et al. reported that components of the nuclear transport machinery were selectively essential for the survival of KRAS-mutant cells, despite their phenotypic heterogeneity [13]. In our previous study, we demonstrated that targeting nuclear export protein XPO1 with selinexor resulted in a robust synthetic lethal interaction with oncogenic KRAS both *in vitro* and *in vivo* [14]. The identification of synthetic lethality between XPO1 and KRAS using computations analysis [14], along with the strong co-expression of these genes observed in PDAC patients from the TCGA dataset, further supports the rationale for co-targeting XPO1 and KRAS as a therapeutic strategy.

It has been reported that the primary mechanism underlying XPO1i sensitivity of KRAS-mutant cancer cells is intolerance to nuclear IκBα accumulation, with consequent inhibition of NF-κB signaling [13]. We have earlier demonstrated that the efficacy of XPO1i selinexor and KRASG12Ci combinations can be mechanistically attributed to the downregulation of NF-κB driven cell survival signaling, as well as induction of cell cycle arrest by reducing CDK4 expression and increasing the nuclear accumulation of tumor suppressor protein RB1 [14]. Similarly, in this study, we observed reduction in cell cycle markers cyclin D1 and CDK4 as well as increased RB1 as the molecular effects of the MRTX1133-Eltanexor combination treatment in PDAC cells.

A recent study by Tripathi et al. has implicated nuclear protein export as an important non-canonical pro-oncogenic function of RAS and reported that mutant KRAS inhibition in lung cancer phenocopies XPO1 inhibition, but the effect of KRAS inhibition on XPO1-dependent activity is independent of canonical KRAS downstream pathways RAF/MEK and PI3K/AKT. It was shown that RAS-GTP complex regulates hydrolysis of Ran-GTP to Ran-GDP, a critical step in XPO1-dependent nuclear export [28]. This regulatory relationship between KRAS and XPO1 lends further credence to our KRASi-XPO1i combination therapy approach for an enhanced antitumor effect. In a separate study, treatment with the XPO1 inhibitor selinexor significantly suppressed tumor growth across ten KRAS-mutant NSCLC PDX models, regardless of the specific KRAS mutation, suggesting a broad dependency of KRAS-mutant cancers on XPO1 [29]. Supporting this, a genome-wide CRISPR/Cas9 screen conducted on 808 cancer cell lines (Cancer Dependency Map Project) identified XPO1 as a dependency in over 90% of lines, classifying it as a ‘common essential gene’ [12]. Additionally, multiple studies have implicated XPO1 as a general vulnerability across multiple cancer types [30–33]. Given that XPO1 is frequently overexpressed in many malignancies [34, 14], it is plausible that XPO1-mediated nuclear export is co-opted by cancer cells as a widespread mechanism contributing to oncogenesis. Therefore, a combination therapy involving XPO1 and KRASG12D inhibitors can be a viable option for recalcitrant PDAC.

It can be speculated that the KRASG12Di-resistant cancer cells would be eradicated by the use of XPO1 inhibitor as a combination partner. This proposition was validated when we generated KRASG12D inhibitor-resistant cancer cell line from KRASG12D-mutant parental AsPC-1 cells and found that this resistant cell line was indeed sensitive to the XPO1i Eltanexor. Our results demonstrate that the combinations of XPO1i with KRASG12Di can effectively inhibit the proliferation of KRASG12D-mutant PDAC cells in 2D and 3D cultures. These combinations have been further shown to suppress the clonogenic potential of KRASG12D-mutant PDAC cells. The combined treatment of MRTX1133 and Eltanexor exhibit pronounced effects on the expression of KRAS downstream signaling molecules and the activity of KRAS associated kinases. Using both ICR-SCID and immunocompetent mice models of KRASG12D-mutant PDAC, we have demonstrated superior efficacy of Eltanexor and MRTX1133 combination in enhancing tumor regression. Moreover, the combination regimen showed appreciable tolerability, causing no significant adverse effects during treatment. Collectively, these findings imply that the inhibition of XPO1 activity could be a plausible therapeutic strategy for overcoming resistance to KRASi as combining Eltanexor with a KRASG12Di can have a durable and synergistic antitumor activity in cancer patients that have developed resistance to therapy with KRASG12Di.

Recently, a phase I/II trial of Exportin 1 inhibitor plus docetaxel in previously treated, advanced KRAS-mutant solid tumors show safety and some encouraging efficacy [35]. This and other trials indicate that Exportin 1 inhibitor may find applicability in KRAS mutant tumor patient population. Results from our study indicate that Eltanexor and MRTX1133 combination can be a viable therapy to effectively suppress KRASG12D-mutant PDAC tumor growth that warrants further investigations [36]. We believe that this preclinical study can make a huge impact toward the development of a KRAS targeted drug combination therapy for the treatment of KRASG12D-mutant PDAC with better therapeutic outcomes.

## Data Availability

The data generated in this study are available on reasonable request from the corresponding author.

## Ethics Declaration

Animal studies were conducted under Wayne State University’s Institutional Animal Care and Use Committee (IACUC) approved protocol (# 22-01-4355) in accordance with the approved guidelines.

## Author Contribution

*Conception* – Asfar S. Azmi, Ramzi M. Mohammad; *Design of the work* – Asfar S. Azmi, Husain Yar Khan, Mohammed Najeeb Al Hallak, Hugo Jimenez, Miguel Tubon, Misako Nagasaka; *Data acquisition* – Husain Yar Khan, Amro Aboukameel, Bin Bao, Ahmet B. Caglayan, Hilmi K. Alkan; *Data analysis* – Husain Yar Khan, Adeeb A. Aboukameel, Md. Hafiz Uddin, Sahar F. Bannoura, Fulya Koksalar Alkan, Gregory Dyson, Yang Shi, Rafic Beydoun; *Interpretation of data* – Asfar S. Azmi, Hasan Korkaya, Allan M. Johansen, Callum McGrath, Grayson Barker, Khalil Choucair, Muhammad Wasif Saif, Anthony F. Shields; *Drafted and revised the manuscript* – Husain Yar Khan, Azeddine Atfi, Philip A. Philip, Bassel El-Rayes, Boris C. Pasche, Asfar S. Azmi

