## Supplemental Figures and Tables for "Co-targeting KRAS and Exportin1 as an effective therapeutic strategy for KRASG12D mutant pancreatic ductal adenocarcinoma"

### Figure S1.

**A.** **HPAF-II**

CI for experimental values:

| MRTX1133 (nM) | KPT8602 (nM) | Fa | CI |
|---|---|---|---|
| 125 | 125 | 0.897751 | **0.131** |
| 250 | 250 | 0.899346 | **0.250** |
| 500 | 500 | 0.913224 | **0.314** |

**B.** **HPAC**

CI for experimental values:

| MRTX1133 (nM) | KPT8602 (nM) | Fa | CI |
|---|---|---|---|
| 125 | 125 | 0.585654 | **0.359** |
| 250 | 250 | 0.670939 | **0.126** |
| 500 | 500 | 0.625339 | **0.625** |

**C.** **AsPC-1**

CI for experimental values:

| MRTX1133 (nM) | KPT8602 (nM) | Fa | CI |
|---|---|---|---|
| 125 | 125 | 0.528231 | **0.614** |
| 250 | 250 | 0.580176 | **0.619** |
| 500 | 500 | 0.60527 | **0.904** |

**D.** **Panc-1**

CI for experimental values:

| MRTX1133 (nM) | KPT8602 (nM) | Fa | CI |
|---|---|---|---|
| 125 | 125 | 0.299984 | **0.207** |
| 250 | 250 | 0.3048 | **0.393** |
| 500 | 500 | 0.298897 | **0.836** |

**E.** **6694c2**

CI for experimental values:

| MRTX1133 (nM) | KPT8602 (nM) | Fa | CI |
|---|---|---|---|
| 125 | 125 | 0.17478 | **0.727** |
| 250 | 250 | 0.379555 | **0.729** |
| 500 | 500 | 0.513621 | 1.101 |

**Figure S1: Synergistic inhibitory effects of KRASG12Di-XPO1i treatments on the growth of several PDAC cell lines**. Different dose combinations of MRTX1133 with KPT8602 (Eltanexor) synergistically inhibit the proliferation of KRASG12D mutant PDAC cell lines **(A)** HPAF-II, **(B)** HPAC, **(C)** AsPC-1, **(D)** Panc-1, and **(E)** 6694c2 as indicated by combination index (CI) values less than 1, determined by CalcuSyn**.** All results are expressed as percentage of control ± S.E.M of six replicates.

### Figure S2.

#### A. HPAF-II (3D)

CI for experimental values:

| MRTX1133 (nM) | KPT8602 (nM) | Fa | CI |
|---|---|---|---|
| 31.25 | 31.25 | 0.661561 | **0.383** |
| 62.5 | 62.5 | 0.742375 | **0.382** |
| 125 | 125 | 0.833192 | **0.287** |
| 250 | 250 | 0.858594 | **0.405** |
| 500 | 500 | 0.896835 | **0.427** |
| 1000 | 1000 | 0.893408 | **0.911** |

#### B. HPAC (3D)

CI for experimental values:

| MRTX1133 (nM) | KPT8602 (nM) | Fa | CI |
|---|---|---|---|
| 62.5 | 62.5 | 0.417725 | **0.543** |
| 125 | 125 | 0.642479 | **0.381** |
| 250 | 250 | 0.873315 | **0.166** |

#### C. KPC-313 (3D)

CI for experimental values:

| MRTX1133 (nM) | KPT8602 (nM) | Fa | CI |
|---|---|---|---|
| 31.25 | 31.25 | 0.256513 | **0.337** |
| 62.5 | 62.5 | 0.392396 | **0.198** |
| 125 | 125 | 0.498429 | **0.170** |
| 250 | 250 | 0.558221 | **0.213** |

**Figure S2: KRASG12Di synergizes with XPO1i to inhibit the viability of PDAC cell lines in 3D cultures**. Synergistic inhibitory effects of the combination treatment at multiple doses of MRTX1133 and KPT8602 (Eltanexor) in constant ratio on the viability of **(A)** HPAF-II, **(B)** HPAC, and **(C)** KPC-313 cells in 3D cultures, as demonstrated by CI < 1 determined by CalcuSyn. All results are expressed as percentage of control ± S.E.M of four replicates.

Figure S3.

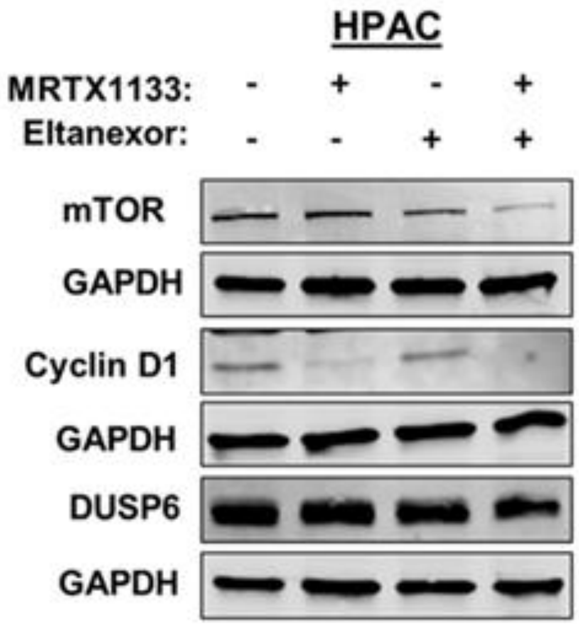


**Figure S3: KRASG12Di-XPO1i modulates KRAS signaling and cell cycle**. Immunoblot showing reduced expression of mTOR, DUSP6 and Cyclin D1 in combination treated HPAC cells treated with 10 nM and 100 nM of MRTX1133 and Eltanexor, respectively.

### Figure S4.

#### Upstream Kinase Analysis (Kinase Score)

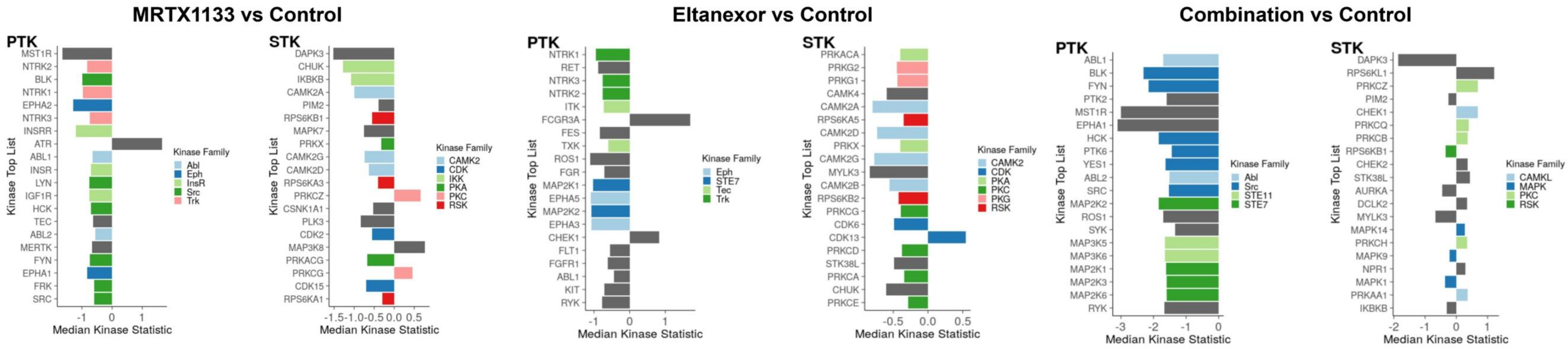


**Figure S4: Upstream Kinase Analysis (UKA).** Following phosphosite analysis of drug treated PDAC cell lysates, UKA was performed using sets of phosphosites, which were linked to each kinase to predict differential kinase activity. Data shows clear reduction in kinase scores of several kinases associated with MAPK signaling in combination treated HPAC cell lysates.

**Figure S5.**

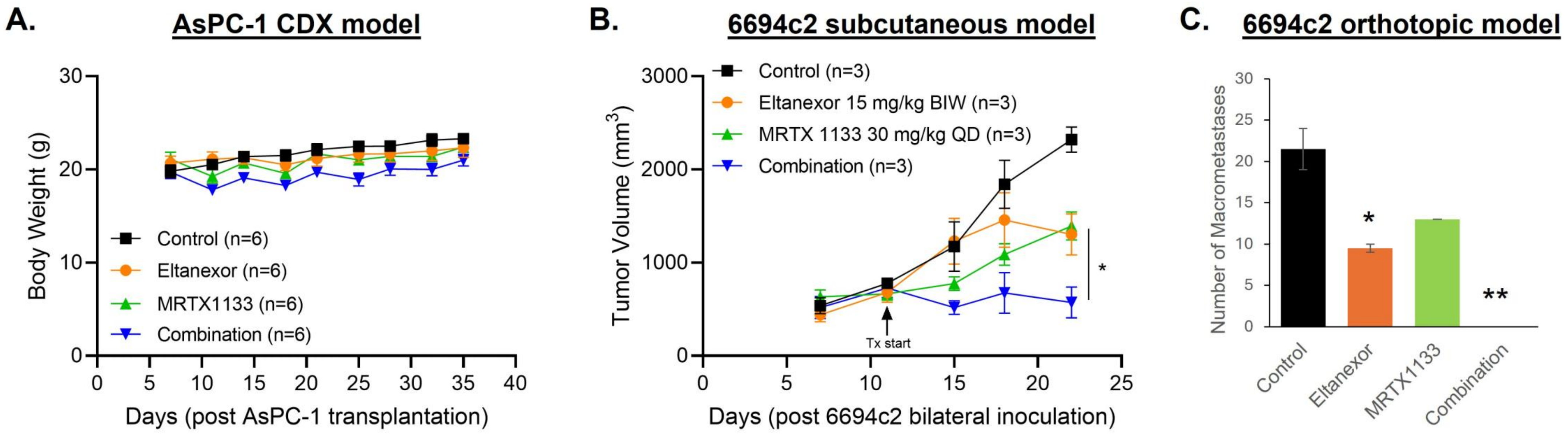


**Figure S5: Antitumor and antimetastatic effects of KRASG12Di-XPO1i combination in PDAC *in vivo* models. (A)** No substantial loss in the animal body weights was observed in the AsPC-1 CDX tumor bearing ICR-SCID mice treated with MRTX1133, Eltanexor or their combination. **(B)** The combination of MRTX1133 and eltanexor demonstrates significantly (* $P < 0.05$) greater suppression of 6694c2 KPC tumor volumes at the end of treatment compared to either of the single agents alone. **(C)** Barplot showing significant loss of total number of macrometastases in combination treated C57BL/6 mice bearing orthotopic 6694c2 allografts. * $P < 0.05$, ** $P < 0.01$ compared to control.

**Supplementary Table 1.** **Sequences of primers used to perform RT-PCR**

| Primers | Sequences | |
|---|---|---|
| MAP2K1 | Forward | CAATGGCGGTGTGGTGTTC |
| | Reverse | GATTGCGGGTTTGATCTCCAG |
| MAP2K2 | Forward | AGGTCCTGCACGAATGCAA |
| | Reverse | CGTCCATGTGTTCCATGCAA |
| MAP2K3 | Forward | GACTCCCGGACCTTCATCAC |
| | Reverse | GGCCCAGTTCTGAGATGGT |
| MAP3K5 | Forward | CTGCATTTTGGGAAACTCGACT |
| | Reverse | AAGGTGGTAAAACAAGGACGG |
| MAP3K6 | Forward | GATGCCTTCTACAACGCGGAT |
| | Reverse | CACGCACACCAAGGTGGTA |
| MST1R | Forward | CTTTGACGTGAAGTACGTGGT |
| | Reverse | CGTATGGCTACAAACACAGCAC |
| ABL1 | Forward | TGAAAAGCTCCGGGTCTTAGG |
| | Reverse | TTGACTGGCGTGATGTAGTTG |
| ABL2 | Forward | GTTGAACCCCAGGCACTAAAT |
| | Reverse | CAACGAAGAGATTAGGGTCACTC |
| ROS1 | Forward | CCACATAATCTGAGTGAACCGTG |
| | Reverse | CGCTGCTACAGCCAACCTC |
| SYK | Forward | CATGGAAAAATCTCTCGGGAAGA |
| | Reverse | GTCGATGCGATAGTGCAGCA |
| β-actin | Forward | GCACAGAGCCTCGCCTT |
| | Reverse | TCATCATCCATGGTGAGCTG |

**Supplementary Table 2.** **Phosphosite analysis.** Number of differentially phosphorylated phosphosites (PTK and STK) between conditions are identified using ANOVA-Dunnett's test.

| Assay Type | PTK | | STK | |
|---|---|---|---|---|
| Comparisons | Up | Down | Up | Down |
| MRTX1133 vs Control* | 4 | 13 | 0 | 4 |
| Eltanexor vs Control* | 1 | 3 | 0 | 1 |
| Combination vs Control* | 0 | 24 | 1 | 6 |
| *Significance was obtained using one-way ANOVA followed by post-hoc Dunnett's test. $p<0.05$ | | | | |
